# Spermidine supplementation accelerates time of pupariation and contributes to early degeneration of silk glands in *Bombyx mori* (Lepidoptera: Bombycidae)

**DOI:** 10.64898/2026.06.15.732520

**Authors:** Brinda Goda Lakshmi Didugu, Anitha Mamillapalli

## Abstract

Spermidine is a ubiquitous biogenic amine that is known to promote growth, development and autophagy. The silk glands of *Bombyx mori* L. (Lepidoptera: Bombycidae) produce mulberry silk, which has high economic value. Silk is released during the formation of a cocoon, and the glands undergo degradation involving both autophagy and apoptosis. The present study explores the effect of spermidine feeding on the maturation of *B. mori* 5^th^ instar larvae with particular emphasis on its role in growth at the end of the larval stage and during the early phase of the silk gland degeneration. 5^th^ instar larvae were divided into two groups and fed with control and spermidine-treated mulberry leaves. At the end of the 5^th^ instar stage, the spermidine group showed a significant increase in body and silk gland weights, which helped in achieving larval critical weight prior to the control group. Significantly elevated levels of *Atg8* expression, an increased number of lysosomes and high chromatin condensation were observed in the Spd group during the early larval – pupal transition phase, which helped in prior silk gland degeneration.

## Introduction

Polyamine, spermidine (Spd), was found to be the most abundant biogenic amine in the anterior, middle and posterior silk glands (SGs) of *Bombyx mori* L. (Lepidoptera: Bombycidae) (*B. mori*) (Hamana et al., 1984). They were shown to play an important role in the growth and development of *B. mori* (Kasa et al., 2023; Lattala et al., 2014; Rajan et al., 2021). Earlier reports showed that polyamine levels in the SGs were found to be maximum during the 5^th^ instar feeding stage and decreased during non-feeding spinning and pre-pupal stages (Didugu & Mamillapalli, 2025). Exogenous supplementation of Spd promoted cell cycle progression in cultured silkworm cells (Chang et al., 2021), and the inhibition of ornithine decarboxylase (ODC), the rate-limiting enzyme in polyamine biosynthesis, by DFMO feeding resulted in decreased polyamine levels and several developmental defects (Rajan et al., 2022).

Several pathways were reported to be involved in the promotion of longevity and autophagy by Spd. It was shown to promote longevity by chromatin remodeling and upregulation of autophagy-related genes in *Drosophila*, *Saccharomyces* and humans (Eisenberg et al., 2009). Spd-induced hypusine was shown to be a crucial indicator for fasting-induced autophagy in yeast, nematodes and human cells (Hofer et al., 2024). It is known as a potential scavenger of reactive oxygen species (ROS), which is crucial for autophagy in mouse fibroblast cells, CHO cells, and HeLa cells (Rider et al., 2007; Scherz-Shouval & Elazar, 2007). Supplementation of Spd enhanced autophagy through specific epigenetic modifications in the honey bee (Kojić et al., 2024).

SGs of *B. mori* produce mulberry silk. 5^th^ instar larvae consume maximum diet, and SGs actively synthesize silk proteins which contribute 30% - 50% to the body weight (Matsuura et al., 1968). Diet was shown to affect larval development, body size, body weight, and timing of metamorphosis in wax moths and honey bees (Mohamed et al., 2014; Nicholls et al., 2021). Larval critical weight (LCW) is the threshold weight required by the larvae to metamorphose into the functional adult. Previous studies from *Drosophila melanogaster* (Almeida de Carvalho & Mirth, 2017; Tyson et al., 2023) and *B. mori* (Keshan et al., 2015; Saha et al., 2009) have shown that the LCW acts as a morphometric indicator to initiate metamorphosis. In silkworms, SGs undergo degeneration during the transition from larvae to the pupal stage. The SG index is also a key indicator to assess the SG growth, silk secretion, and gland degeneration during metamorphosis (Gu et al., 2024; Murugesh et al., 2013). Degeneration of silk glands was shown to involve autophagy and apoptosis during the larval–pupal transition period, exhibiting features of membrane blebbing, initiation of chromatin condensation, increased acid phosphatase and *Atg8* expression levels (Goncu & Parlak, 2008; Li et al., 2011; Montali et al., 2017).

Though the role of Spd in growth and autophagy was studied in different organisms, its effect on *B. mori* larval maturation and SG degeneration was not explored. The present study specifically investigates the cellular events associated with the autophagic phase of silk gland degeneration following Spd supplementation. Larval critical weight, SG index, biochemical and cellular changes were assessed in Spd-supplemented SGs. The results showed that Spd-supplemented larvae achieved LCW earlier and showed accelerated SG degeneration when compared to the control group.

## Materials and Methods

### Silkworm rearing and feeding protocol of Spd

*B. mori* (CSR2×CSR4) larvae were procured during the 4^th^ instar stage from the Andhra Pradesh State Government Sericulture Centre, Chebrolu, East Godavari District, Andhra Pradesh, India. Rearing was performed under the standard temperature of 28 ± 2 °C and 80 – 85 % of relative humidity till complete cocoon formation. The start day of the 5^th^ instar stage after the 4^th^ moult is considered as day 0 (D0). Twenty-four hours after feeding from D0 is considered as day 1 (D1). D7, D9, and D11 correspond to the last day of the 5^th^ instar feeding stage, spinning, and prepupal stages, respectively. Larvae were divided into control (Con) and Spd groups. For each rearing, three experimental replicates were maintained per group, with 25 larvae in each replicate (n = 150). A total of two rearings were conducted for the analysis. The Con group was fed with V1 variety mulberry leaves, and the Spd group was fed with leaves supplemented with 50 µM Spd (RM 5438; HiMedia Laboratories, India). The concentration of Spd was based on the standardization of the feeding experiment in the previous study (Lattala et al., 2014). The detailed experimental plan of the study is given in Fig. 1.

**Fig 1.**
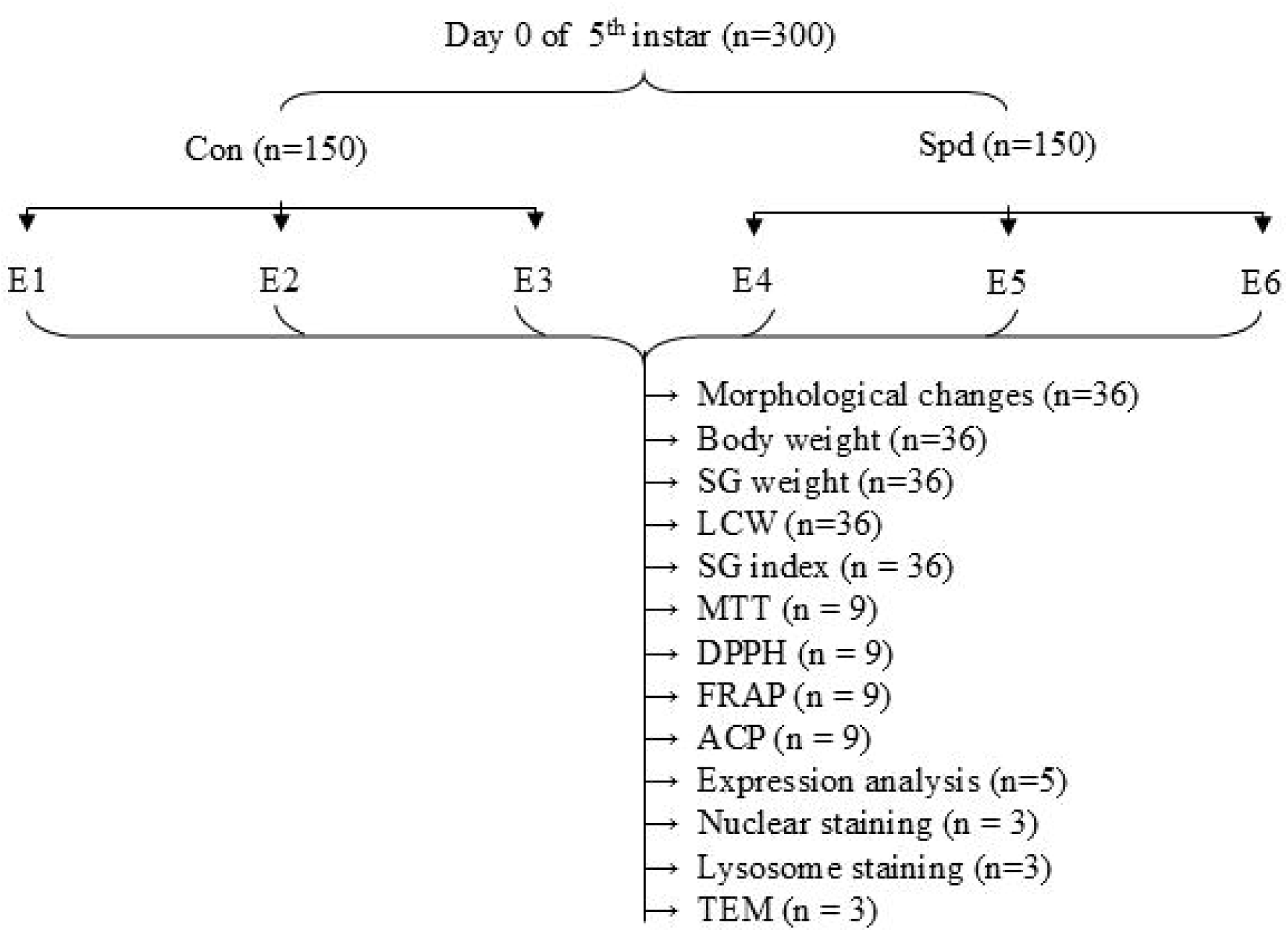
Experimental plan used in the study. E1 to E6 represent experimental replicates of the Con and Spd groups (n = 50 in each replicate).

### Assessment of morphological changes from D7 to D11 in the Con and Spd groups

Larvae/cocoons were picked up randomly on D7, D9, and D11 from both Con and Spd groups for assessment of the morphological changes and maturation status (n = 36) during larval–prepupal transition.

### Determination of larval critical weight in Con and Spd groups

Larval weights were determined by recording the individual body weights from the Con and Spd groups (n = 36) as per the published method. Measurements were taken every 24 hours from D7 (the last feeding day) to check maturation differences between the Con and Spd groups. Larval critical weight (LCW) refers to the threshold body weight that the larvae must attain to enter the pupal stage. Bilinear regression is a statistical method used to fit two linear relationships to larval growth data, where the intersection point represents the critical weight at which further development becomes independent of nutritional input. The mathematical model used in this study was obtained from a previously published study in *Drosophila* (Tyson et al., 2023) using the pwlf library in Python (code deposited in GitHub https://github.com/AnithaMamillapalli/Silk-gland-autophagy/blob/438c216941120501089a8f2877dd845ea856bb8c/bilr.ipynb).

### Assessment of the silk gland weights and silk gland index of the Con and Spd groups

To determine the SG weight and index, larvae were randomly selected on D7, D9, and D11 from the Con and Spd groups (n = 36). Individual SG weights were measured using an electronic weighing balance. The SG index was determined on D7, D9, and D11as described earlier (Gu et al., 2024). The equation used in the analysis is given below.

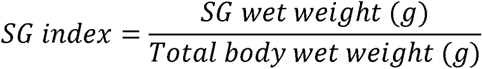

### Evaluation of the metabolic activity in the silk gland of the Con and Spd groups

Metabolic activity of SGs was determined by MTT assay (3-(4,5-Dimethylthiazol-2-yl)-2,5-diphenyltetrazolium bromide) as per the published protocol (Nath et al., 2005; Yerra et al., 2016) (n = 9). SGs were isolated from the Con and Spd groups on D7, D9, and D11. SGs isolated from each larva were incubated for 1 hour in 100 µL of MTT (5mg/ml) and 900 µL of PBS. MTT was removed, and the glands were washed with PBS. The formazan formed was solubilized with overnight incubation in 1 mL of 50 % Tween 80. The solubilized formazan was collected in a separate microfuge tube and diluted with PBS (1:1, v/v), and the optical density was recorded at 570nm using a Shimadzu UV spectrophotometer. The values obtained were reported as absorbance (OD units) per gland.

### Assessment of antioxidant potential in the silk gland of the Con and Spd groups

Antioxidant potential was determined in Con and Spd groups of SGs on D7, D9, and D11 by 2,2,-Diphenyl-1-Picrylhydrazyl (DPPH), and Ferric Reducing Antioxidant Power (FRAP) (n=9) as per the established protocols (Yerra et al., 2016).

Total SG was homogenized in 1mL of 0.1M phosphate buffer of pH 7.0. To 200 µl of the SG homogenate, 800 µL of 0.1 mM DPPH was added, and the mixture was incubated in the dark for 30 min. The absorbance was recorded at 520nm.

To the 200 µL of SG homogenates, 2.8 mL of FRAP reagent (300 mM sodium acetate buffer, pH 3.6, 10 mM TPTZ, 20 mM FeCl_3_) was added and incubated at 37 °C for 30 min. The absorbance readings were recorded at 593nm. Both DPPH and FRAP results were extrapolated to total SG and plotted.

### Quantification of acid phosphatase activity in the silk glands of the Con and Spd groups

Lysosomal enzyme activity of SGs in Con and Spd groups on D7, D9, and D11 was determined by acid phosphatase activity (ACP) (n=9) according to a published protocol (Montali et al., 2017). Different concentrations (units) of the commercially available ACP (P3752; Merck) were used to make the standard plot. SGs were homogenised in 1ml of 0.1M phosphate buffer, and the supernatant was used for the assay. To 75 µL of the supernatant, 500 µL of 0.1 M citrate buffer, 400 µL of 15 mM para-nitrophenyl phosphate (PNPP), and 525 µL of distilled water were added, and the mixture was incubated at 37 °C in a water bath for 30 minutes. Reaction was terminated by the addition of 2.5mL of 0.2 N NaOH (28 °C for 15 minutes), and absorbance was recorded at 410 nm. The protein content of the SGs was determined by the Lowry method (Lowry et al., 1951). ACP Enzyme activity is expressed in units per mg of protein.

### Gene expression analysis in the silk gland of the Con and Spd groups by RT-qPCR

Total RNA was isolated from the entire SGs of the D7, D9, and D11 using the TRIzol method (n =5). RNA quantity and quality were determined using a nanodrop. RNA (2.5 µg) was converted into cDNA using RT-PCR Kit (iScript™ cDNA Synthesis Kit, Bio-Rad). qPCR was performed in a 10 µl reaction volume, containing 5 µl of 2X SYBR Green Mix (Sso

Advanced Universal SYBR® Green Supermix), 2 µl of cDNA (1:10 dilution), 6.6 pm of each primer and 1 µL of autoclaved water. The primer sequences of *Atg8* and *Rp49* were obtained from previously published literature and are mentioned in Supplementary Table 1. The relative expression of the *Atg8* gene was calculated using *Rp49* as a reference gene and expressed as 2^-ΔΔCt^ values. Negative controls (no template and no reverse transcriptase) were also carried out to check for contamination of genomic DNA and non-specific amplification.

### Sectioning of PSG for staining

SGs were isolated from Con and Spd larvae on D7, D9, and D11. PSGs were separated and fixed in 4% formaldehyde at 4 °C for 24 h (n=3), then washed in PBS. PSGs from three larvae were pooled for paraffin block preparation, and cross-sections were obtained at a thickness of 3 µm using a microtome. The slides were dried in a hot air oven prior to staining.

### Assessment of lysosomes with Lysosight green

The sections were stained with Lysosight green (FSB-004-2024, Fluoresight bioprobes private limited) for 15 minutes and were washed with PBS for 5 minutes, followed by DAPI staining. The stained cross-sections were observed under a confocal microscope (Olympus FV 4000) to assess the lysosome abundance. Images obtained at different magnifications were processed using cellSens (Olympus), ZEISS ZEN 3.9 (ZEISS), and ImageJ software.

### Assessment of nuclear morphology of PSG cross sections from Con and Spd groups with DAPI and Hoechst staining

Nuclear staining of PSG cross sections was performed with 4′,6-diamidino-2-phenylindole (DAPI) (5 µg for 5 minutes) (MB097; HiMedia Laboratories, India) and Hoechst 33258 (10 ng for 10 minutes) (H6024; Sigma-Aldrich, USA) separately. The stained cross-sections were observed under a confocal microscope (Olympus FV 4000) to assess the changes in nuclear morphology. Images obtained at different magnifications were processed using cellSens (Olympus), ZEISS ZEN 3.9 (ZEISS), and ImageJ software. From each slide, three cross-sections were randomly selected for edge detection, chromatin condensation, perimeter and area analyses of PSG sections in both Con and Spd groups. Chromatin condensation was checked with the intensity of DAPI and Hoechst staining on D7, D9, and D11. The images are processed as grayscale images by the OpenCV library and processed as unsigned 8-bit integers. Staining of chromatin in the PSG section on D7 stage from the Con group, with low levels of chromatin condensation information, was used as a baseline for assessing the chromatin in D9 and D11. The baseline was obtained by the mean of the D7 stage of the Con group, by excluding pixels with zeros. With this basis, all further images were scaled to ensure only pixels greater than the threshold showed color, while the others were set to zero. To ensure consistent colouring across a single replicate, the maximum intensity was obtained from the D11 of the Spd group. To ensure appropriate linear representation of coloring, we utilized the colorcets library of colormaps (Kovesi, 2015). This process ensures a visual linear representation of the amount of chromatin above a certain threshold across a single replicate. (Code is deposited in GitHub https://github.com/AnithaMamillapalli/Silk-gland-autophagy/blob/438c216941120501089a8f2877dd845ea856bb8c/intensity.ipynb).

### Edge detection and evaluation of the perimeter and area of PSG sections in Con and Spd groups

The DAPI-stained PSG sections on D7, D9, and D11 of Con and Spd groups were analysed for geometric parameters of edge detection, perimeter, and area (n = 3). The parameters were selected based on the morphological changes the SGs undergo during degeneration, namely, membrane invaginations and shrinkage of area. The images, as unsigned 8-bit integers, were processed using the OpenCV library for edge detection. To ensure the edge was appropriately identified, all pixels with a value above a certain noise threshold were set to 255. The noise parameter was tuned to be the lowest value to produce an acceptable edge detection in relation to the original figure. This binary image made up of completely black and white pixels was then passed through the Sobel edge detection algorithm. Some images (D7 and D9 of the Spd group) were then passed through a morphological closing process to ensure uniform and consistent edges. And finally, the edge was extracted, and analysis on perimeter and area was possible though built-in OpenCV functions. The noise parameter and optional morphological closing create a subjective approach for each image, but the parameters for Sobel edge detection and other operations remained uniform throughout. This approach ensures high-quality edges for all images. Finally, the pixel-based measurements of area and perimeter were appropriately scaled to ensure comparisons were accurate. (Calculation codes are deposited on GitHub https://github.com/AnithaMamillapalli/Silk-gland-autophagy/blob/438c216941120501089a8f2877dd845ea856bb8c/edge%20detection.ipynb).

### Ultrastructural analysis of PSGs using transmission electron microscopy of the Con and Spd groups

Ultrastructural imaging of PSGs from isolated SGs was carried out on the D9 (n = 3). PSGs were immediately fixed in 4 % glutaraldehyde and stored at 4 °C. The samples were processed according to a previously published protocol (Chunduri et al., 2022). PSG sections were stained with lead citrate and uranyl acetate and observed under transmission electron microscopy. TEM imaging was performed using a JEOL JEM-2100 electron microscope.

### Statistical analysis of the data

All data were presented as mean ± standard error of the mean (SEM). All graphs were constructed using GraphPad Prism 8.0 software. The results were statistically analyzed using a two-way analysis of variance (ANOVA) to evaluate the effects of treatment during different developmental days when compared to the Con. Post hoc multiple comparisons using Tukey’s honest significant difference (HSD) test were used to check significance levels between the groups. ***, **, and * were used for the statistical significance *p* < 0.001, *p* < 0.01, and *p* < 0.05 levels respectively.

## Results

### Spd feeding promotes early maturation of B. mori larvae

Nutrition and growth help in the proper development of silkworms, which determines the 5^th^ instar larval duration in *B. mori*. In the present study, the effect of Spd supplementation on the maturation of larvae to the pupal stage was checked. Two groups were maintained, and the feeding was given as detailed in the methodology section (Fig. 1). Con and Spd groups were evaluated for morphological changes, body weights, and SG weights on D7, D9, and D11. Larvae in the Spd group were found to undergo maturation to the prepupal stage earlier than the Con group (Fig. 2A). Both the Con and Spd groups showed a highly significant decrease in body weight from D7 to D9 and from D9 to D11, respectively. A highly significant increase in body weight was observed on D7 in the Spd group when compared to the Con group (Fig. 2B). No significant difference in body weight was observed between the Spd and Con groups on D9 and D11. As the decision to pupate depends on larval critical weight, bi-segmental linear regression was performed with body weight and time of pupariation in both the Con and Spd groups. Results showed that the Spd group took 32.5 hours from D7 to attain LCW, while the Con group took 43.6 hours (Fig. 2C). Therefore, the 5^th^ instar larvae of the Spd group entered the pupal stage 11.1 hours earlier when compared to the Con group.

**Fig. 2.**
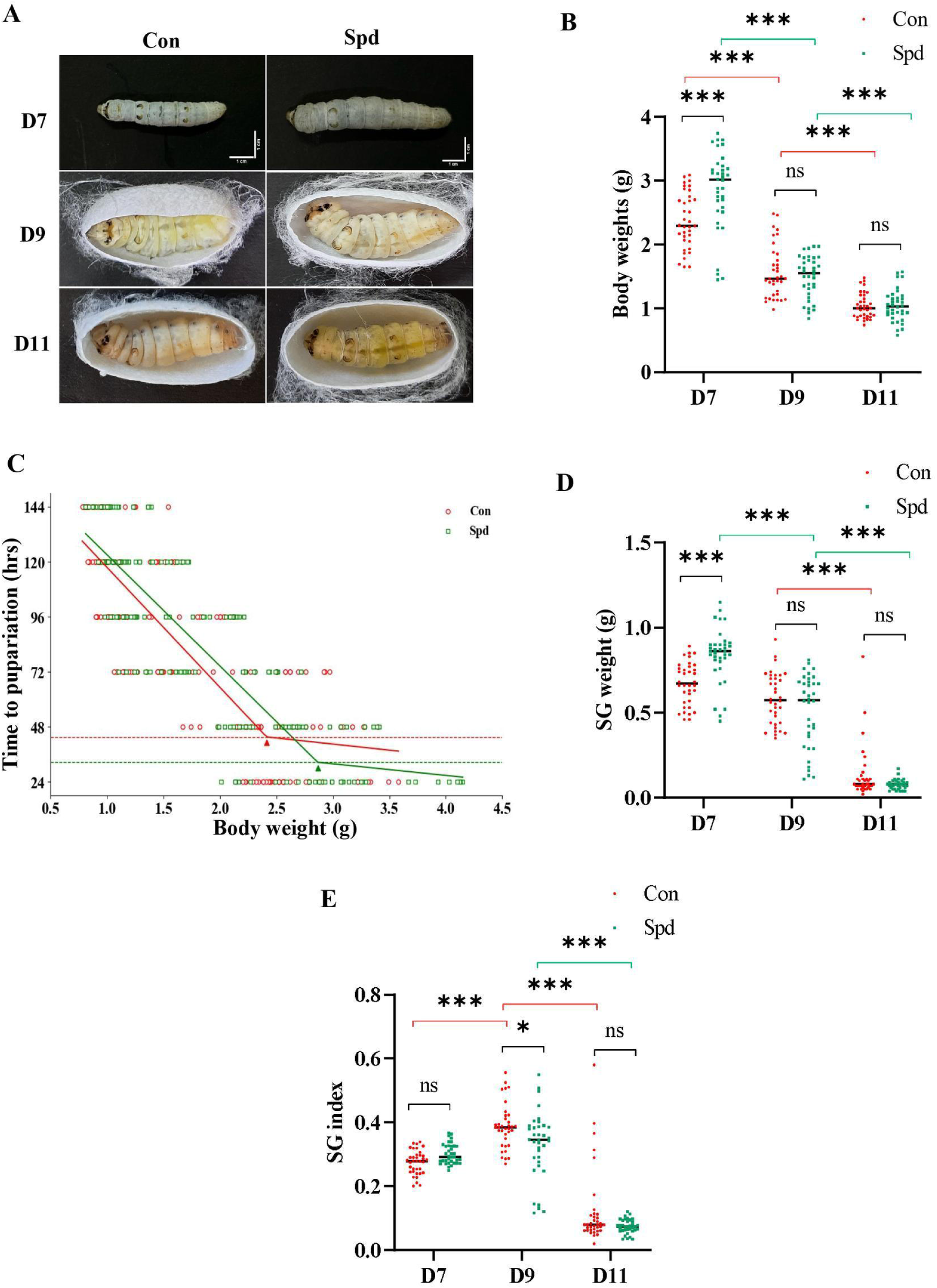
Effect of Spd on the development of *B. mori* larvae. The parameters (A) morphological changes, (B) larval body weight, (C) time to pupariation (TTP), (D) SG weight, and (E) SG index of Con and Spd groups during D7, D9 and D11 were assessed (n=36). \*\*\**p* < 0.001, \*\**p* < 0.01, \**p* < 0.05. Black, red, and green lines indicate statistical significance between Con vs. Spd, Con vs Con, and Spd vs Spd, respectively.

In the 5^th^ instar larval stage, SG is the major contributor to body weight. SG weights were measured in both groups on D7, D9, and D11. In the Con group, SG weights showed a highly significant decrease from D9 to D11. The Spd group showed a significant decrease in SG weight from D7 to D9 and from D9 to D11. The Spd group showed a highly significant increase in SG weight on D7 when compared to the Con group. No significant change in the SG weight was observed in the Spd group when compared to the Con group on D9 and D11 (Fig. 2D). The significant increase in SG weights in the Spd group compared to the Con group on D7 could have helped larval maturation earlier in the treated group.

The SG index was assessed to evaluate the gland degeneration in the Con and Spd groups on D7, D9, and D11. The Con group showed a highly significant increase in the SG index from D7 to D9. Both the Con and Spd groups showed a highly significant decrease in the SG index from D9 to D11. The SG index did not show any significant change on D7 between the Con and Spd groups. The Spd group showed a significant decrease in SG index when compared to the Con group on D9, which could have promoted early degradation of SG in the Spd group when compared to the Con group (Fig. 2E, Supplementary Table 2).

### Exogenous Spd supplementation accelerates the degeneration of SGs

MTT assay measures the metabolic activity of the cells, and the degeneration of SGs was checked by the assay on D7, D9 and D11. There was no significant difference in the metabolic activity in the Spd group when compared to the Con group across D7, D9, and D11. Both groups showed a highly significant decrease in metabolic activity from D7 to D9 (Fig. 3A). No significant decrease in metabolic activity was observed from D9 to D11 in both the groups. DPPH and FRAP assays revealed that there was no significant difference in the antioxidant activity in the Spd group when compared to the Con group across D7, D9, and D11 (Fig. 3B & C, Supplementary Table 3).

**Fig. 3.**
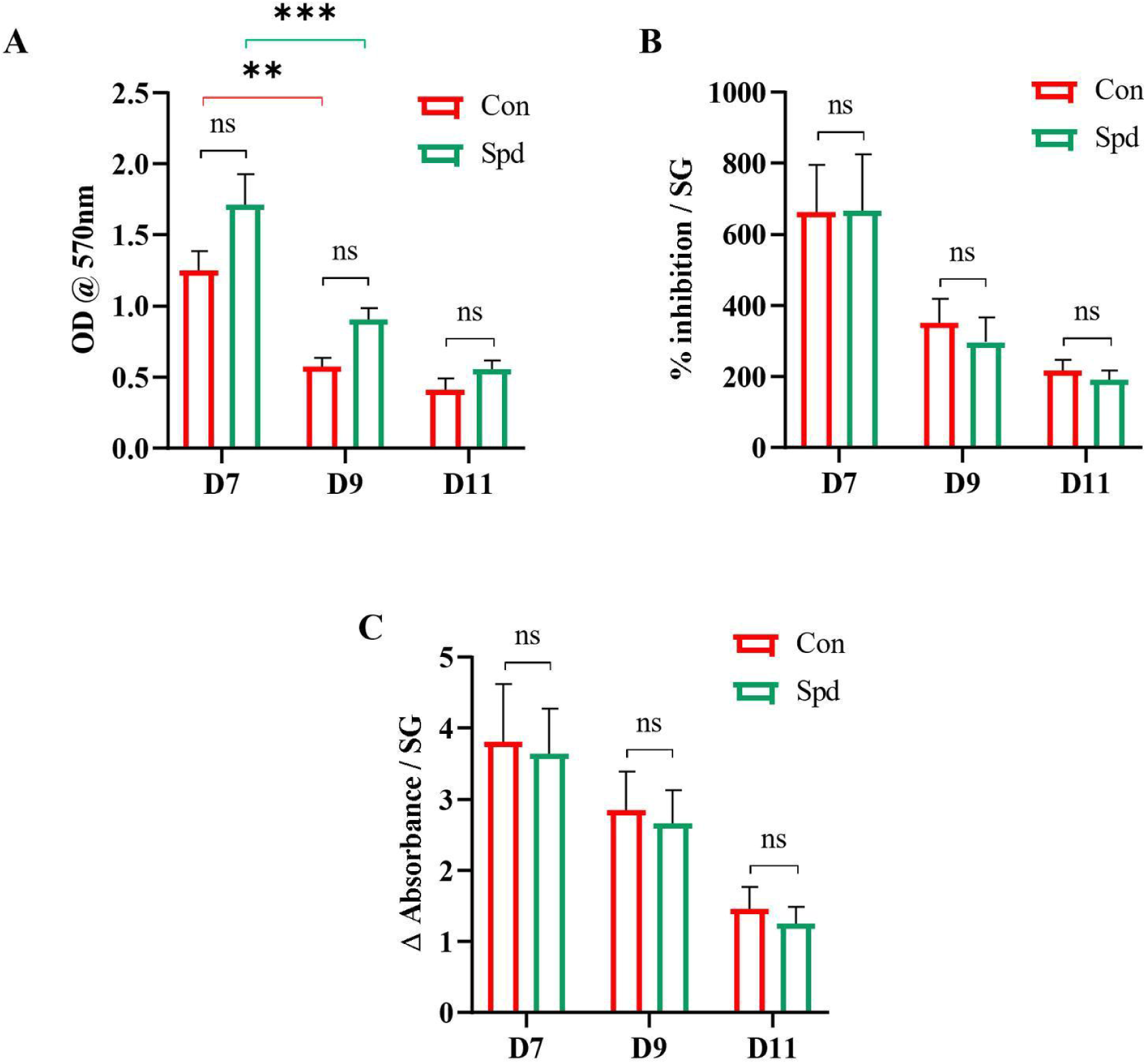
Effect of Spd on biochemical parameters of SG on D7, D9, and D11. (A) MTT assay, (B) DPPH assay, (C) FRAP, in Con and Spd groups (n=9). \*\*\**p* < 0.001, \*\**p* < 0.01, \**p* < 0.05. Black, red, and green lines indicate statistical significance between Con vs. Spd, Con vs Con, and Spd vs Spd, respectively.

### Spd promotes autophagy of SGs

The degeneration of SGs includes both autophagy and apoptosis. Based on the role of Spd on autophagy, the markers of autophagy were assessed in Con and Spd glands on D7 and D9.

ACP is a lysosomal marker of autophagy, and its activity was measured in SGs of Con and Spd groups on D7 and D9. Spd group showed an increase in ACP activity on D7 when compared to the Con group. Both the Con and Spd groups showed an increase in activity from D7 to D9 (Fig. 4A, Supplementary Table 4).

**Fig. 4.**
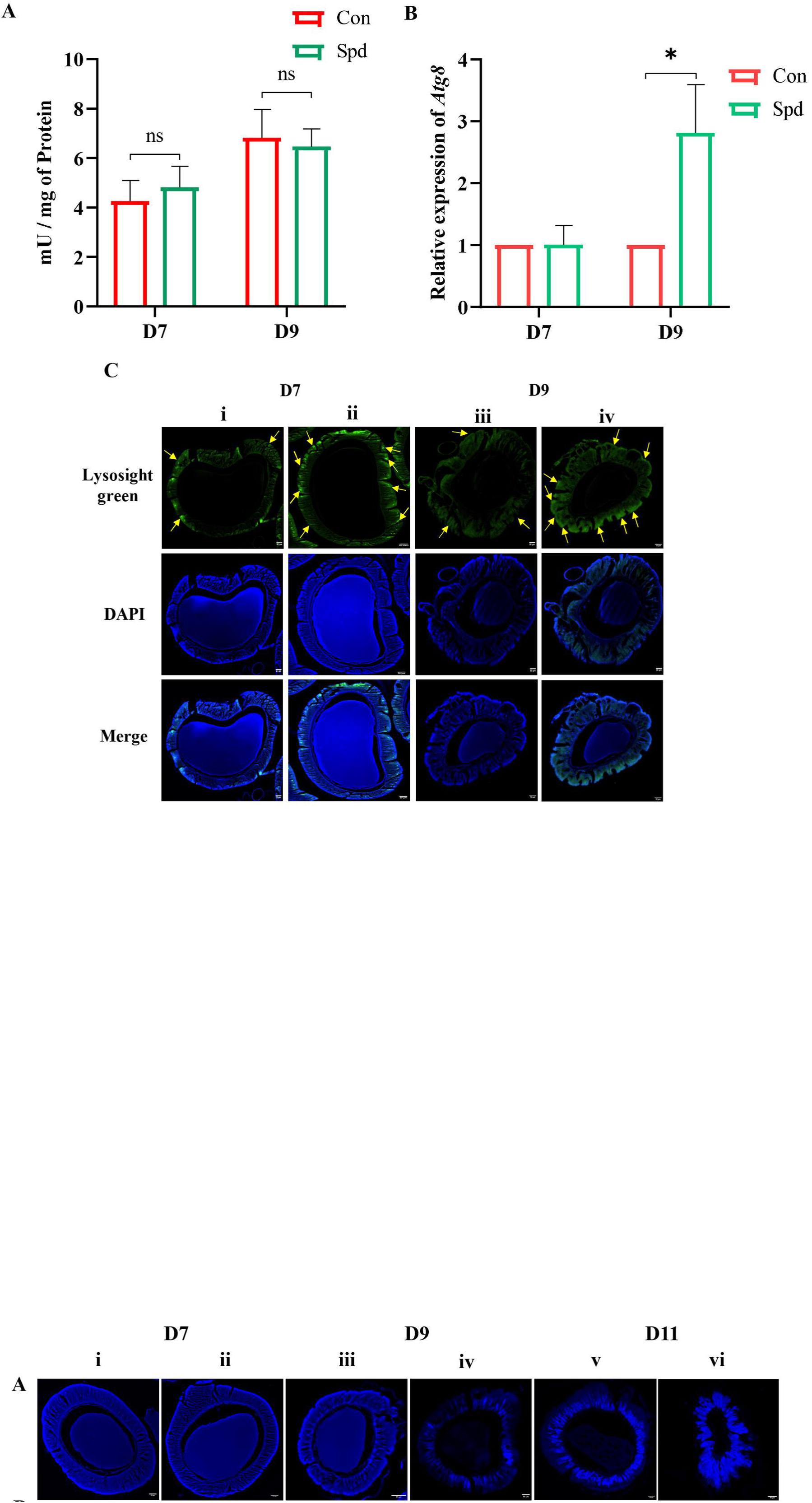
Assessment of Spd supplementation on autophagy of SGs on D7 and D9. (A) ACP activity (n=9), (B) relative expression of *BmAtg8* expression, (n=5) and (C) lysosomal staining of PSG section (n=3) (Con- i and iii, Spd- ii and iv). Black, red, and green lines indicate statistical significance between Con vs. Spd, Con vs Con, and Spd vs Spd, respectively (\**p* < 0.05). Yellow arrows represent lysosomes in different panels of C. The scale bar in each panel denotes 50 µM.

*Atg8* is a crucial gene for the closure of the autophagosome during the autophagy process of SGs in *B. mori*. Thus, RNA expression analysis of *Atg8* was carried out in the SGs of the Con and Spd groups on D7 and D9. There is no significant change in the *Atg8* expression in the Spd group on D7 when compared to the Con group. The Spd group reported a significant increase in fold change in the *Atg8* expression on D9 when compared to the Con group (Fig. 4B, Supplementary Table 4).

Visualization of lysosomes in the PSG cross sections of both groups was performed by staining with lysosight green. PSGs of the Spd group showed a high intensity of staining of lysosomes when compared to the Con group on both D7 and D9 (Fig. 4C). Thus, ACP activity, *Atg8* expression and lysosomal staining showed Spd supplementation promoted autophagy of SGs.

### Spd facilitates SG degeneration by enhancing chromatin condensation

SGs undergo multiple rounds of endoreplication and show highly branched and dispersed chromatin. The branched chromatin undergoes condensation during SG degradation. As Spd binds to the negatively charged DNA and promotes chromatin condensation, the changes in the chromatin were checked in both the Con and Spd groups on D7, D9, and D11 by DAPI staining. Results showed that the chromatin was uniformly spread and branched throughout the PSG on D7 in both Con and Spd groups (Fig. 5A i and ii). Chromatin condensation was observed on D9 (Fig. 5A iii and iv), which significantly increased by D11 in the Spd group when compared to the Con group (Fig. 5A v and vi).

**Fig. 5.**
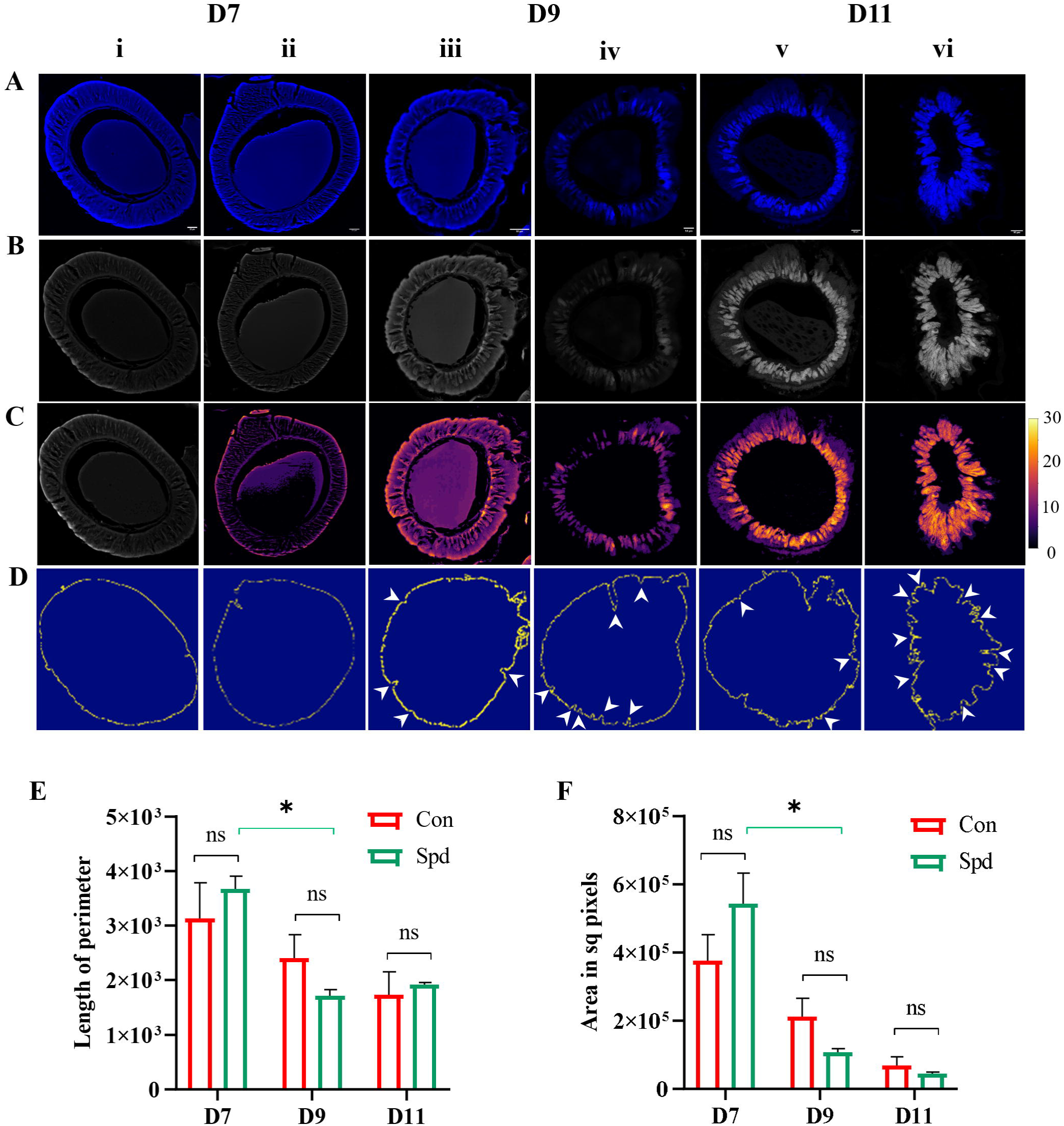
Morphology and chromatin changes of PSG cross sections on D7, D9, and D11 in Con and Spd groups. (A) DAPI staining, (B) Gray scale images of DAPI, (C) Heatmap images generated from DAPI staining, (D) Edge detection of PSG, (Con- i, iii and v, Spd- ii, iv and vi) (E) Perimeter of PSG cross-section, and (F) Area of PSG sections during D7, D9 and D11 (n=3). The white arrowheads indicate invagination of the epithelium. The scale bar in each panel denotes 50 µM. Black, red, and green lines indicate statistical significance between Con vs. Spd, Con vs Con, and Spd vs Spd (\**p* < 0.05).

Chromatin condensation was further studied by heatmap analysis as described in the methodology section. Results showed that the Spd group showed an increase in chromatin condensed on D7, D9, and D11 when compared to the Con group (Fig. 5B and 5C). PSGs stained with Hoechst correlated with DAPI staining (Fig. S1).

Membrane invaginations lead to blebbing, which is associated with autophagy. The outline of the PSG membrane was constructed as detailed in the methods section. Invaginations of the PSG epithelium were observed more prominently in the Spd group on D9 and D11 (Fig.5D iv and vi) when compared to the Con (Fig. 5D iii and v).

Analysis of the perimeter and area of PSG cross sections was performed to assess the shrinkage quantitatively. The Con group did not show a significant decrease in the perimeter of the PSG cross sections from D7 to D9 and from D9 to D11. Spd group showed a significant decrease in the perimeter of the PSG cross sections from D7 to D9 and no significant difference from D9 to D11. Spd group showed an increase in the perimeter when compared to the Con group on D7 (Fig. 5E).

Analysis of the area of PSG cross sections did not show a significant decrease from D7 to D9 and from D9 to D11 in the Con group. Spd group showed a significant decrease in the area of the PSG cross sections from D7 to D9 and no significant difference from D9 to D11. Spd group showed an increase in the area when compared to the Con group on D7 (Fig. 5F, Supplementary Table 5).

These results show that Spd supplementation contributed to the increased chromatin condensation and shrinkage of PSGs, which could have led to early degradation of SGs.

### Ultrastructural analysis of PSG sections reveals autophagy structures in the Spd group

Ultrastructural analysis was carried out to check for the autophagy-related organelles in PSG sections of the Con and Spd groups on D9. Analysis revealed the presence of early endosomes in the Con PSG sections and late endosomes in the Spd PSG sections (Fig. 6A). PSGs of the Spd group showed mature autophagosomes (Fig. 6B). PSGs of the Spd group showed highly condensed chromatin when compared to the Con group, which reiterated the DAPI results (Fig. 6C). These results show that the PSGs of the Spd group show signs of early degeneration.

**Fig. 6.**
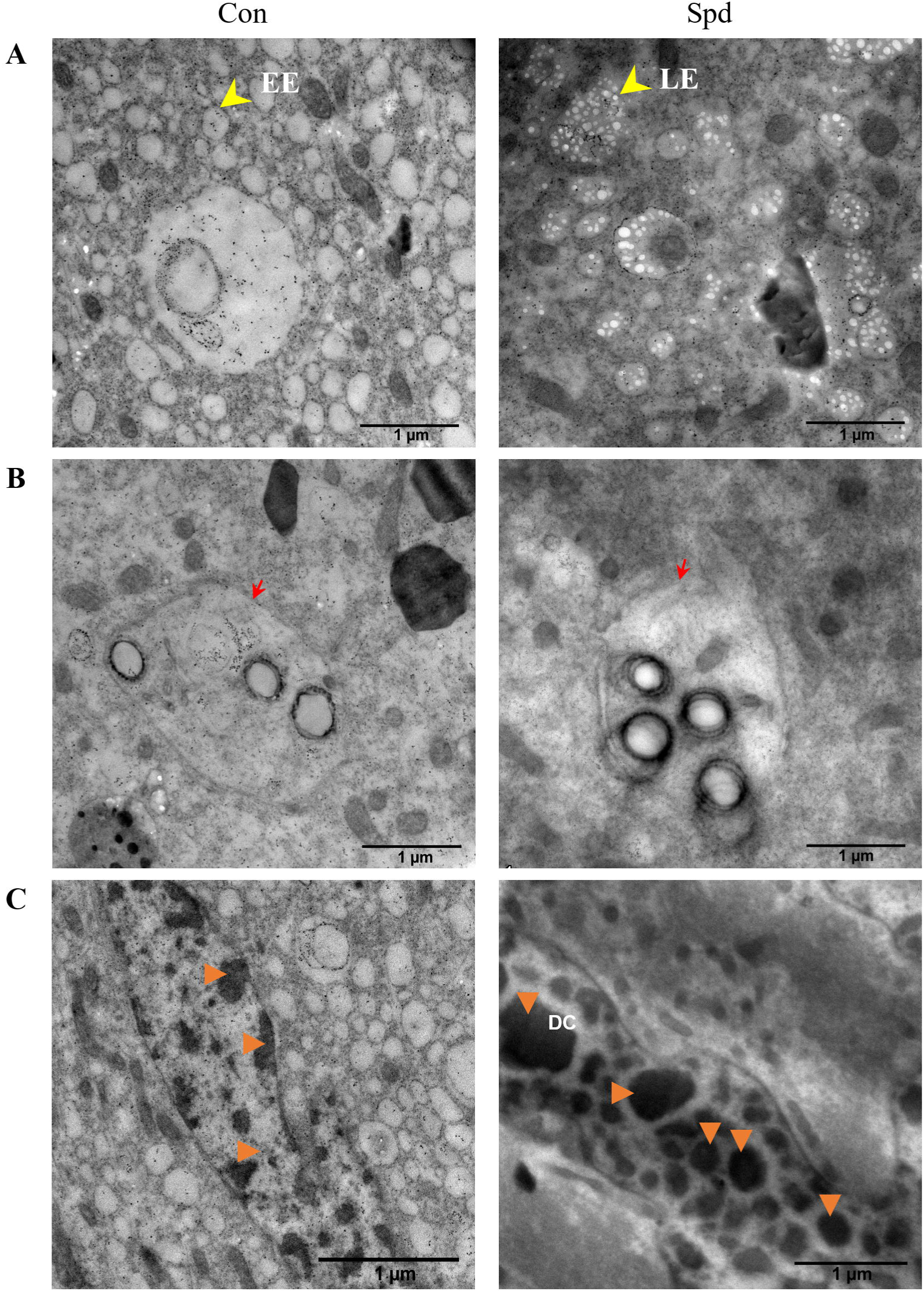
Ultrastructural analysis of PSG cross sections of Con and Spd on D9 stage. (A) TEM images of PSG cross sections showing early endosomes (EE), and late endosomes (LE) (yellow arrow heads), (B) PSG cross sections showing autophagosomes (red arrow heads), (C) SG showing darkly stained condensed chromatin (orange arrow heads) in Con and Spd groups. The scale bar represents 1 µM.

**Fig. 7.**
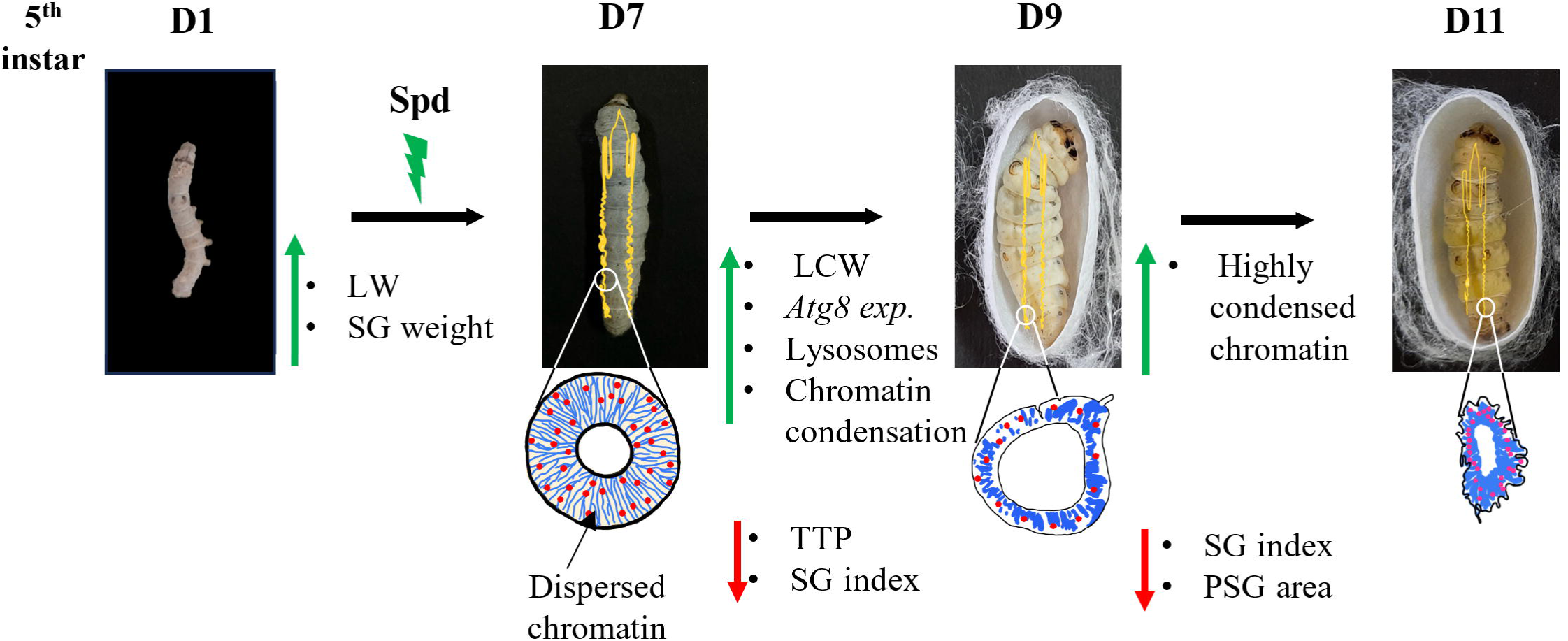
Schematic representation of Spd-mediated maturation of *B. mori* larvae led to the early degeneration of SGs. Green arrows indicate an increase, red arrows indicate a decrease, blue lines represent chromatin in SG sections, and red dots represent Spd. LW – Larval weight, SG – Silk gland, LCW – Larval critical weight, TTP – Time to pupariation, PSG-Posterior silk gland.

## Discussion

Exogenous Spd supplementation was shown to play a crucial role in the growth and development of several organisms, such as *Rattus norvegicus* (Jeevanandam et al., 1997), *Fusarium graminearum* (Tang et al., 2021), including *B. mori* (Lattala et al., 2014). The present study showed that supplementation of Spd to 5^th^ instar *B. mori* larvae showed a significant increase in the body weight and SG weight on D7, which could have led the larvae to reach critical weight before the Con group. The result coincides with the studies which showed that diet affects larval development, body size, body weight, and timing of metamorphosis in honeybees, wax moths and *Drosophila* (Almeida de Carvalho & Mirth, 2017; Mohamed et al., 2014.; Nicholls et al., 2021; Saha et al., 2009; Tyson et al., 2023). The silk gland index represents the relative weight of the silk gland to the total body weight and is used to assess growth and developmental changes. The present study showed that the Spd group showed a significant decrease in the SG index, a sign of SG degeneration (Gu et al., 2024). Results from the present study reveal that the Spd group attained body weight and entered the pre - pupal stage earlier than the Con group.

SG is the largest organ in the 5^th^ instar larval stage, responsible for the synthesis of the silk proteins and is metabolically highly active. The current study assessed the viability of the SG by MTT assay, and the Spd group showed an increase in the viability, but did not find any significant difference between the Con and Spd groups. A study conducted on SH-SY5Y cell lines with different concentrations of Spd observed a significant increase with 0.1 and 1µM concentration (Fairley et al., 2023). It could be possible that many concentrations of Spd were not checked in the current study to find a significant difference. Spd supplementation in honey bees reduced oxidative stress and increased antioxidant potential (Đorđievski et al., 2023). Spd induces autophagy and reduces oxidative stress in granulosa cells of Sichuan white geese (Jiang et al., 2023). SGs of the Con and Spd groups were processed to evaluate the antioxidant potential. It was observed that the antioxidant potential significantly decreased from D7 to D9 and D11, but no significant difference was observed in the Spd group when compared to the Con group. In our previous studies, Spd feeding was found to significantly enhance metabolic activity and antioxidant potential during the 5th instar larval stage (Yerra et al., 2016). Therefore, the reasons for the results obtained in the present study could be due to the days selected or could also be due to the single concentration of Spd used in the study.

The role of Spd in autophagy of SGs was assessed by ACP activity, *Atg8* expression and lysosomal staining. The increase in ACP activity observed in the present study correlated with the earlier studies, which showed its association with the degeneration of SGs (Ghonmode & Tembhare, 2007; Goncu & Parlak, 2008). SGs of the Spd group showed a significant upregulation of *Atg8* on D9 when compared to the Con group. The *Atg8* expression was shown to be upregulated in the anterior silk glands of *B. mori,* showing autophagy-related SG degeneration during the prepupal stage (Li et al., 2011). A similar result of *Atg8* expression was observed in SG after Spd supplementation. PSG cross sections stained with lysosight green also revealed an increase in lysosomal abundance in the Spd group when compared to the Con group. Thus, Spd supplementation enhanced the autophagy markers and contributed to early degradation of SGs.

The larval-pupal transition in *B. mori* involves reorganization of chromatin in the SG. The posterior silk gland (PSG) cells exit from the endomitotic cycle before the anterior and middle SG cells during the late 5^th^ instar larval stage (Zhang et al., 2012). The Spd group showed highly condensed chromatin on D9 and D11 when compared to the Con group, indicating an accelerated progression of cellular remodeling as reported earlier (Li et al., 2010). The result is similar to the binding studies of Spd with genomic B-DNA of the chromatin (Deng et al., 2000). Membrane invaginations observed in SG epithelium at the D9 and D11 stages in the current study were consistent with those reported in previous studies, which showed invaginations as a hallmark of SG degeneration (Matsuura et al., 1968). The presence of late endosomes and mature autophagosomes by TEM analysis in the SGs of the Spd group confirmed their early degradation, as reported earlier (Li et al., 2010; Neikirk et al., 2023).

The present study concludes that Spd supplementation to the 5th instar *B. mori* larvae increased body and SG weights with a reduction in the time to pupariation. Reduced silk gland index, increased chromatin condensation, and shrinkage of SGs could have promoted early degradation of SGs. The significant increase in *Atg8* expression could have contributed to the early degeneration of SGs in the Spd group. A more detailed study on the influence of the Spd on the SG transcriptome needs to be carried out.

## Acknowledgements

The authors would like to thank the Andhra Pradesh State Government Sericulture Centre, Chebrolu, for providing silkworms; MURTI facility, GITAM for real-time PCR, confocal microscopy; GIMSR for providing the microtome; and CCMB for the TEM facility. The authors wish to thank Krishna Calindi (MS Computer Science student at Texas A&M, College Station, Texas, USA) for writing Python programs for the critical weight, Correlation analysis, edge detection, and heatmaps of the DAPI and Hoechst staining analyses. B.G.L.D. wishes to thank the University Grant Commission, Ministry of Education, Government of India, for Ph.D. scholarship (UGCES-22-GE-AND-F-SJSGC-15816).

## Conflicts of interest

The authors declare that there are no competing interests.

## Author contributions

B.G.L.D.: Investigation, Formal analysis, and writing - original draft. A.M.: Conceptualization, Supervision, Writing - Review, and Editing.

## Funding

No funding.

## Data availability

The data supporting the findings of this study are available from the corresponding author (A.M.) upon request. Python codes are available on GitHub: https://github.com/AnithaMamillapalli/Silk-gland-autophagy.

## Supplementary Figure legends

**Fig. S1.** Hoechst staining of the PSGs cross sections on D7, D9, and D11 in Con and Spd groups. (A) Hoechst staining, (B) Heatmap images of Con and Spd groups during larval-prepupal transition. The scale bar in each panel denotes 50 µM.

## Supplementary Tables

**Supplementary Table 1.** Details of the primers used for the amplification of *Atg8* and *Rp49* genes from SGs of *B. mori*.

**Supplementary Table 2.** Results of the Tukey Multiple Comparison Test of body weights, SG weights, and SG index of Con and Spd groups on D7, D9 and D11.

**Supplementary Table 3.** Tukey Multiple Comparison Test on the biochemical parameters of *B. mori* SGs following Spd supplementation on D7, D9 and D11.

**Supplementary Table 4.** Results of the Tukey Multiple Comparison Test on ACP activity and *Atg8* expression in SGs of *B. mori* following Spd supplementation on D7 and D9.

**Supplementary Table 5.** Results of the Tukey Multiple Comparison Test on perimeter and area of PSG sections of *B. mori* following Spd supplementation on D7, D9, and D11.

## Notes

### Competing Interest Statement

The authors have declared no competing interest.

