## Supplementary figures and images for "Spermidine supplementation accelerates time of pupariation and contributes to early degeneration of silk glands in *Bombyx mori* (Lepidoptera: Bombycidae)"

### Supplemental Figure 1

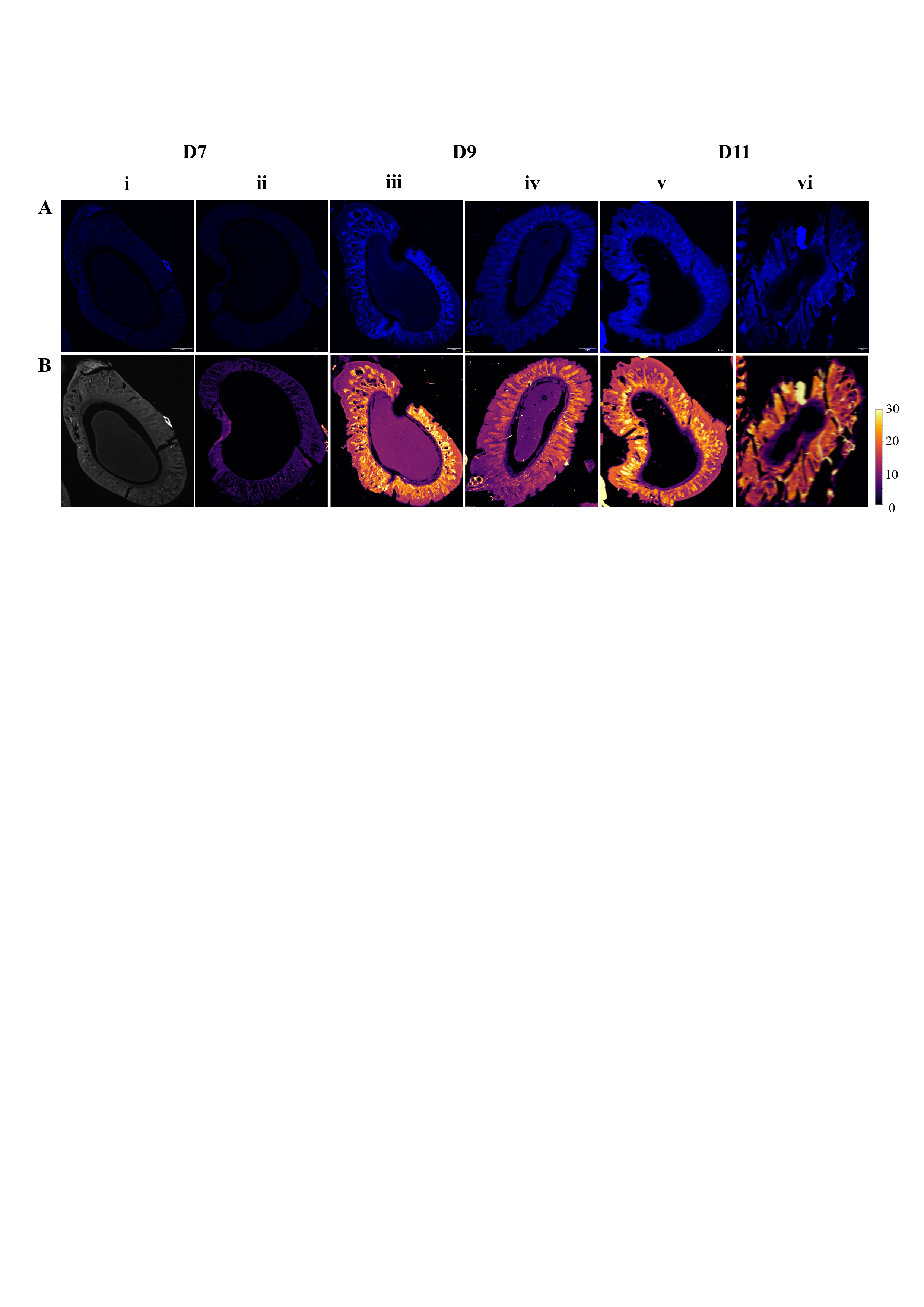
