## Supplemental Table 1 for "Spermidine supplementation accelerates time of pupariation and contributes to early degeneration of silk glands in *Bombyx mori* (Lepidoptera: Bombycidae)"

Supplementary Table 1. Details of the primers used for the amplification of *Atg8* and *Rp49* genes from SGs of *B. mori*.

| **S. No.** | **Primer Name** | **Sequence (5'→3')** | **Product size** | **Reference** |
| --- | --- | --- | --- | --- |
| 1 | *Atg8-*F | CCAGATCGCGTTCCTGTAAT | 205 | (Montali et al., 2017) |
| 2 | *Atg8-*R | GAGACCCCATTGTTGCAGAT |  |  |
| 3 | *Rp49-*F | CAGGCGGTTCAAGGGTCAATAC | 213 | (Liu et al., 2016) |
| 4 | *Rp49-*R | TGCTGGGCTCTTTCCACGA |  |  |
