## Supplemental Table 2 for "Spermidine supplementation accelerates time of pupariation and contributes to early degeneration of silk glands in *Bombyx mori* (Lepidoptera: Bombycidae)"

Supplementary Table 2. Results of the Tukey Multiple Comparison Test of body weights, SG weights, and SG index of Con and Spd groups on D7, D9 and D11.

| **Tukey's multiple comparisons test** | | | | | | |
| --- | --- | --- | --- | --- | --- | --- |
|  | **Body weights** | | **SG weights** | | **SG index** | |
|  | **Significant** | **Adjusted *P Value*** | **Significant** | **Adjusted *P Value*** | **Significant** | **Adjusted *P Value*** |
| **D7:Con vs. D7:Spd** | *** | <0.001 | *** | <0.001 | ns | 0.6491 |
| **D7:Con vs. D9:Con** | *** | <0.001 | ns | 0.1355 | *** | <0.001 |
| **D7:Con vs. D9:Spd** | *** | <0.001 | *** | <0.001 | * | 0.0198 |
| **D7:Con vs. D11:Con** | *** | <0.001 | *** | <0.001 | *** | <0.001 |
| **D7:Con vs. D11:Spd** | *** | <0.001 | *** | <0.001 | *** | <0.001 |
| **D7:Spd vs. D9:Con** | *** | <0.001 | *** | <0.001 | *** | <0.001 |
| **D7:Spd vs. D9:Spd** | *** | <0.001 | *** | <0.001 | ns | 0.5529 |
| **D7:Spd vs. D11:Con** | *** | <0.001 | *** | <0.001 | *** | <0.001 |
| **D7:Spd vs. D11:Spd** | *** | <0.001 | *** | <0.001 | *** | <0.001 |
| **D9:Con vs. D9:Spd** | ns | 0.967 | ns | 0.3071 | * | 0.0156 |
| **D9:Con vs. D11:Con** | *** | <0.001 | *** | <0.001 | *** | <0.001 |
| **D9:Con vs. D11:Spd** | *** | <0.001 | *** | <0.001 | *** | <0.001 |
| **D9:Spd vs. D11:Con** | *** | <0.001 | *** | <0.001 | *** | <0.001 |
| **D9:Spd vs. D11:Spd** | *** | <0.001 | *** | <0.001 | *** | <0.001 |
| **D11:Con vs. D11:Spd** | ns | >0.9999 | ns | 0.6358 | ns | 0.0654 |
