## Supplemental Table 3 for "Spermidine supplementation accelerates time of pupariation and contributes to early degeneration of silk glands in *Bombyx mori* (Lepidoptera: Bombycidae)"

Supplementary Table 3. Tukey Multiple Comparison Test on the biochemical parameters of SG of *B. mori* following Spd supplementation on D7, D9 and D11.

| **Tukey's multiple comparisons test** | | | | | | |
| --- | --- | --- | --- | --- | --- | --- |
|  | **MTT** |  | **DPPH** | | **FRAP** | |
|  | **Significant** | **Adjusted *P Value*** | **Significant** | **Adjusted *P Value*** | **Significant** | **Adjusted *P Value*** |
| **D7:Con vs. D7:Spd** | ns | 0.083 | ns | >0.9999 | ns | >0.9999 |
| **D7:Con vs. D9:Con** | ** | 0.0027 | ns | 0.2009 | ns | 0.799 |
| **D7:Con vs. D9:Spd** | ns | 0.3326 | ns | 0.0825 | ns | 0.655 |
| **D7:Con vs. D11:Con** | *** | 0.0001 | * | 0.0182 | * | 0.0367 |
| **D7:Con vs. D11:Spd** | ** | 0.0019 | * | 0.0108 | * | 0.0179 |
| **D7:Spd vs. D9:Con** | **** | <0.0001 | ns | 0.1908 | ns | 0.9 |
| **D7:Spd vs. D9:Spd** | *** | 0.0002 | ns | 0.0776 | ns | 0.7881 |
| **D7:Spd vs. D11:Con** | **** | <0.0001 | * | 0.017 | ns | 0.064 |
| **D7:Spd vs. D11:Spd** | **** | <0.0001 | * | 0.01 | * | 0.0327 |
| **D9:Con vs. D9:Spd** | ns | 0.3752 | ns | 0.9982 | ns | 0.9999 |
| **D9:Con vs. D11:Con** | ns | 0.9275 | ns | 0.908 | ns | 0.4628 |
| **D9:Con vs. D11:Spd** | ns | >0.9999 | ns | 0.8291 | ns | 0.3089 |
| **D9:Spd vs. D11:Con** | ns | 0.0548 | ns | 0.9906 | ns | 0.6191 |
| **D9:Spd vs. D11:Spd** | ns | 0.3162 | ns | 0.9682 | ns | 0.4475 |
| **D11:Con vs. D11:Spd** | ns | 0.9556 | ns | >0.9999 | ns | 0.9998 |
