## Supplemental Table 4 for "Spermidine supplementation accelerates time of pupariation and contributes to early degeneration of silk glands in *Bombyx mori* (Lepidoptera: Bombycidae)"

Supplementary Table 4. Results of the Tukey Multiple Comparison Test on ACP activity and *Atg8* expression in SGs of *B. mori* following Spd supplementation on D7 and D9.

| **Tukey's multiple comparisons test** | | | | |
| --- | --- | --- | --- | --- |
|  | **ACP** | | ***Atg8*** | |
|  | **Significant** | **Adjusted *P Value*** | **Significant** | **Adjusted *P Value*** |
| **D7:Con vs. D7:Spd** | ns | 0.9717 | ns | >0.9999 |
| **D7:Con vs. D9:Con** | ns | 0.2087 | ns | >0.9999 |
| **D7:Con vs. D9:Spd** | ns | 0.3302 | * | 0.0327 |
| **D7:Spd vs. D9:Con** | ns | 0.4125 | ns | >0.9999 |
| **D7:Spd vs. D9:Spd** | ns | 0.5798 | * | 0.0337 |
| **D9:Con vs. D9:Spd** | ns | 0.9921 | * | 0.0327 |
