## Supplemental Table 5 for "Spermidine supplementation accelerates time of pupariation and contributes to early degeneration of silk glands in *Bombyx mori* (Lepidoptera: Bombycidae)"

Supplementary Table 5. Results of the Tukey Multiple Comparison Test on perimeter and area of PSG sections of *B. mori* following Spd supplementation on D7, D9, and D11.

| **Tukey's multiple comparisons test** | | | | |
| --- | --- | --- | --- | --- |
|  | **Perimeter** | | **Area** | |
|  | **Significant** | **Adjusted *P Value*** | **Significant** | **Adjusted *P Value*** |
| **D7:Con vs. D7:Spd** | ns | 0.9042 | ns | 0.3077 |
| **D7:Con vs. D9:Con** | ns | 0.7458 | ns | 0.3206 |
| **D7:Con vs. D9:Spd** | ns | 0.1568 | * | 0.0385 |
| **D7:Con vs. D11:Con** | ns | 0.1638 | * | 0.0161 |
| **D7:Con vs. D11:Spd** | ns | 0.2781 | ** | 0.0096 |
| **D7:Spd vs. D9:Con** | ns | 0.2369 | ** | 0.0093 |
| **D7:Spd vs. D9:Spd** | * | 0.0296 | ** | 0.0011 |
| **D7:Spd vs. D11:Con** | * | 0.0311 | *** | <0.001 |
| **D7:Spd vs. D11:Spd** | ns | 0.0569 | *** | <0.001 |
| **D9:Con vs. D9:Spd** | ns | 0.7849 | ns | 0.7526 |
| **D9:Con vs. D11:Con** | ns | 0.7992 | ns | 0.464 |
| **D9:Con vs. D11:Spd** | ns | 0.9388 | ns | 0.3157 |
| **D9:Spd vs. D11:Con** | ns | >0.9999 | ns | 0.9945 |
| **D9:Spd vs. D11:Spd** | ns | 0.9985 | ns | 0.9577 |
| **D11:Con vs. D11:Spd** | ns | 0.999 | ns | 0.9995 |
